# Energetics of the Transmembrane Peptide Sorting by Hydrophobic Mismatch

**DOI:** 10.1101/2024.02.02.578561

**Authors:** Balázs Fábián, Matti Javanainen

**Affiliations:** Department of Theoretical Biophysics, MPI Biophysics, DE-60438, Frankfurt am Main, Germany; Institute of Biotechnology, University of Helsinki, FI-00790 Helsinki, Finland

## Abstract

Hydrophobic mismatch between a lipid membrane and embedded transmembrane peptides or proteins plays a role in their lateral localization and function. Earlier studies have resolved numerous mechanisms through which the peptides and membrane proteins adapt to mismatch, yet the energetics of lateral sorting due to hydrophobic mismatch has remained elusive due to the lack of suitable computational or experimental protocols. Here, we pioneer a molecular dynamics simulation approach to study the sorting of peptides along a membrane thickness gradient. Peptides of different lengths tilt and diffuse along the membrane to eliminate mismatch with a rate directly proportional to the magnitude of mismatch. We extract the 2-dimensional free energy profiles as a function of local thickness and peptide orientation, revealing the relative contributions of sorting and tilting, and suggesting their thermally accessible regimes. Our approach can readily be applied to study other membrane systems of biological interest where hydrophobic mismatch, or membrane thickness in general, plays a role.

**TOC Graphic:** 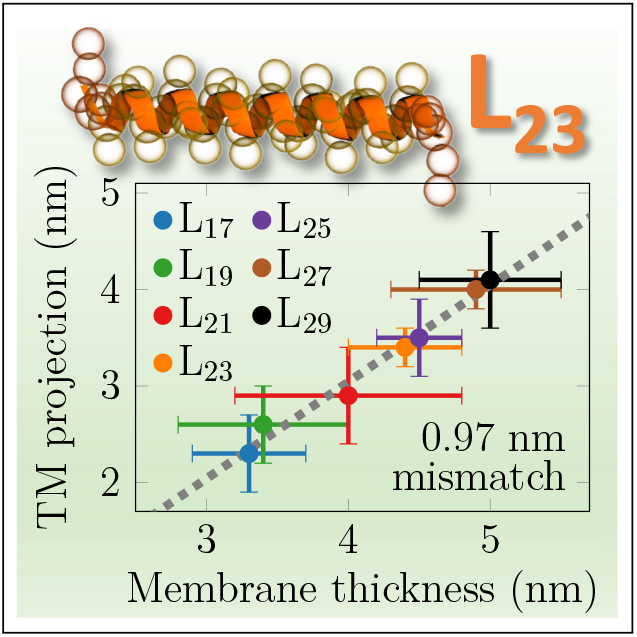

---

Cellular membranes display varied lipid compositions and hence physico-chemical properties supporting the processes taking place in and on them. ^1^ The membranes of the endoplasmic reticulum, the Golgi apparatus, and the plasma membrane differ in their thicknesses,^2^ suggesting that hydrophobic mismatch (MM) controls the sorting and targeting of transmembrane (TM) peptides and proteins along the secretory pathway. MM is the difference in the hydrophobic extents of the transmembrane domain (TMD) and the host membrane^3,4^ which—if unaccounted for—leads to an energetic penalty. Apart from organelle-level sorting, MM also affects protein function, conformation, stability, orientation, oligomerization, and dynamics.^5–8^ Moreover, the plasma membrane^9^ and organelle membranes^10^ are heterogeneous with different local lipid pools manifested in different membrane properties—including thickness—leading to lateral sorting of proteins.^11^ Sometimes, MM cannot be eliminated by lateral sorting, but instead either the membrane, the protein, or both adapt to eliminate the mistmatch. For multi-pass TMDs, this is not straightforward without conformational changes. However, single-pass TMDs can cope with mismatch by multiple mechanisms.

In the case of positive MM (TMD longer than membrane thickness), the TMD can either tilt or bend,^5,12–15^ and the membrane can also respond by locally adjusting its thickness. ^16,17^ This mechanism is intriguing, since it can drive TMD aggregation by lowering the total energetic penalty associated with membrane deformation.^14^ In the case of negative MM, the membrane can bulge, possibly coupled with TMD aggregation.^13,16,17^ Alternatively, water can penetrate the headgroup region to hydrate charged protein residues.^13^ In extreme MM scenarios, the TMDs might alternate their anchoring between membrane leaflets, or even abandon their TM orientation.^15,16^ Of exceptional interest is the related energetics, as it maps the TMD and membrane properties to the preferred response. For example, a single-pass TMD with MM*>*0 might partition to a thicker domain or stay put and increase its tilt. Both actions would at least partially eliminate MM, but which one is favoured? Or are both active within the limits of the thermal energy? Surprisingly, these questions have escaped earlier computational and experimental examinations, likely due to methodological challenges.

Here, we tackle these questions using coarse-grained (CG) molecular dynamics (MD) simulations. We set up a lipid membrane containing a thickness gradient by merging four single-component membranes made of different phosphatidylcholine (PC) lipids with increasing acyl chain lengths: DYPC (3 coarse-grained beads per chain), DOPC (4 beads), DGPC (5 beads) and DNPC (6 beads), corresponding to 12–24 carbon atoms per chain in the Martini 2 model^18,19^ (Fig. 1A). The thicknesses of corresponding single-lipid membranes—ranging from 3.34 to 4.95 nm—are indicated by markers in Fig. 1C. To generate a constant thickness profile along the *x* axis, the different lipid types are maintained at certain *x*-coordinate intervals using flat-bottom potentials (Fig. 1B). A smooth profile in the *x* coordinate range of 8–32 nm is generated with an optimized amount of lipid type overlap (see methods and Fig. S2 in the Supporting Information (SI)), resulting in maximum thickness difference of ≈1.85 nm.

**Figure 1:**
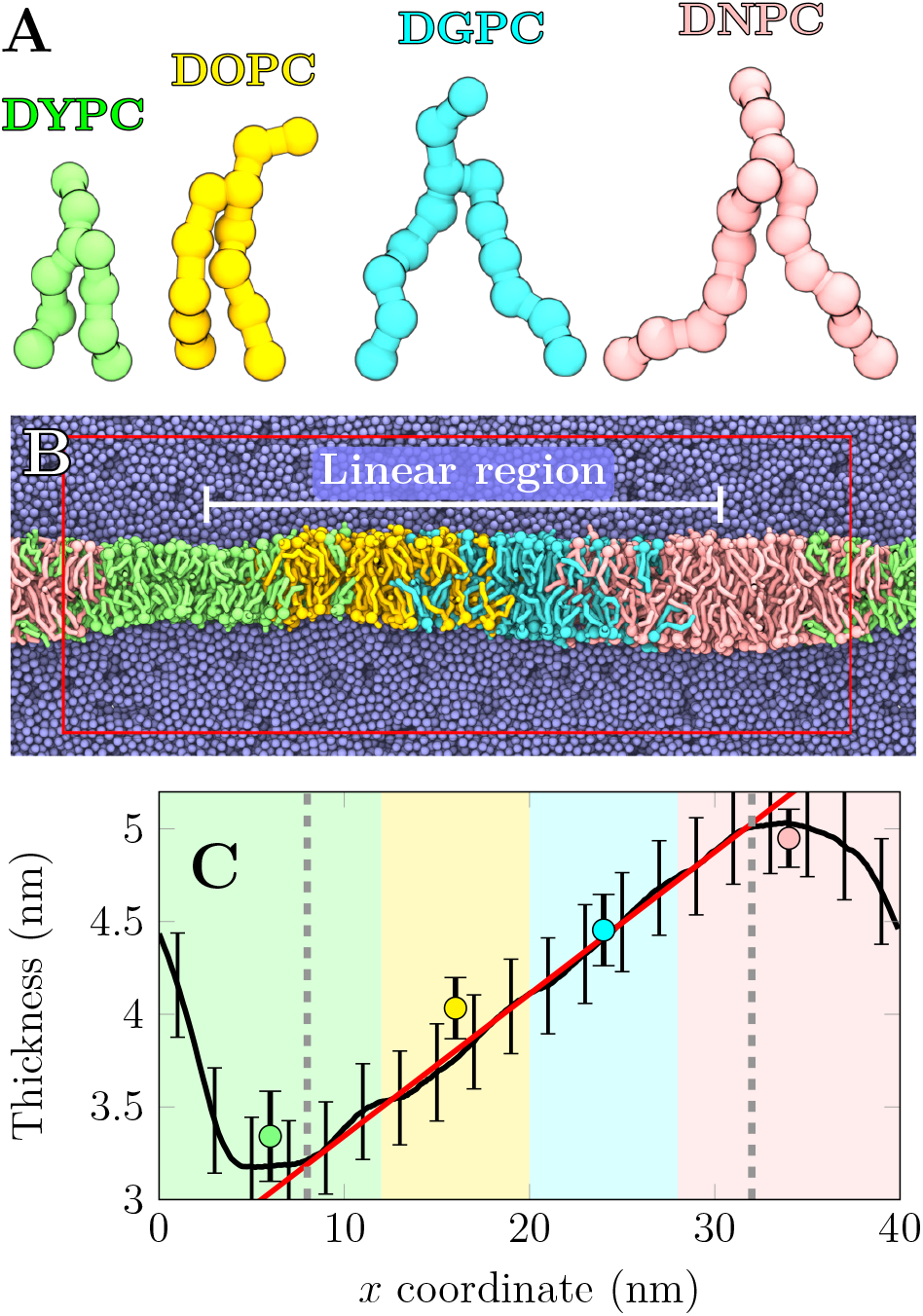
Thickness gradient generation. **A)** The four lipid types used in this study (see SI). Naming follows the Martini convention (`cgmartini.nl`). **B)** Membrane after 5 μs of simulation. Colouring as in A). Water in iceblue. The red rectangle shows the unit cell and white line the linear region (see panel C). **C)** Local membrane thickness (black). A linear thickness gradient with a slope of 0.077 nm/nm (red) is present in between gray dashed lines. Markers show single-component membrane thicknesses at the *x* coordinate in the middle of the corresponding patch in the assembled membrane. Shaded areas show positioning of membrane patches (overlap omitted).

Our approach extends the toolkit available for the studies of hydrophobic mismatch and overcomes some limitations of earlier works. Our setup contains a smooth thickness gradient instead of a two-phase membrane arrangement.^20,21^ Moreover, by using different lipid types to generate the gradient, our approach allows to study membranes in their native, tensionless state and hence with realistic lateral lipid densities.^22^ Unlike with the setup in Ref. 22, a standard GROMACS version is suitable to run the simulations.

Due to periodic boundary conditions, there is an abrupt thickness jump at the edge of the simulation box, but analyses here focus on the linear region. The shape of the flat-bottom potentials is provided in Fig. S1C, whereas Fig. S1E shows the resulting spatial distribution of the different lipid types. The local area per lipid values, shown in Fig. S1D, are in reasonable agreement with values from single-component membrane simulations. Details of system setup and simulations are available in the SI.

We first studied whether the thickness gradient would sort lipids, as highlighted by similar earlier works.^20,22^ To this end, we released the flat-bottom restraints for 5 lipids of each type in both leaflets, and allowed them to sample the membrane freely during a 50 μs simulation. Curiously, the sorting tendency turned out to be minor with free energy differences of ≈1 *k*_B_*T* (Fig. S1F), yet the analysis also revealed a small repulsive bias of similar magnitude (≈1 *k*_B_*T*) at the mixed lipid regions despite our attempts to optimize the overlap of the lipid patches (Fig. S2).

We next embedded polyleucines (Leu17–Leu29) whose hydrophobic thicknesses *l*_TM_ ranged from 2.55 to 4.35 nm to this membrane (Table 1). Polyleucines have demonstrated tolerance for large MM and maintain their TM orientation.^23^ The peptides were capped by two lysines to anchor them to the membrane–water interfaces.^24^ See SI for further details.

**Table 1:** Peptides used in this study. “*l*_TM_” is the length of the hydrophobic region (0.15 nm per leucine). Equilibrium values are provided for local thickness (*d*_eq_), peptide tilt angle (*θ*_eq_), and projected peptide length (*l*_proj_), and *l*_mis_ is the calculated MM.

| Peptide properties |  |  | Unbiased simulations |  |  |  | US simulations |  |
| --- | --- | --- | --- | --- | --- | --- | --- | --- |
| Name | Sequence | $l_{\text{TM}}$ | $d_{\text{eq}}$ (nm) | $\theta_{\text{eq}}$ (°) | $l_{\text{proj}}$ (nm) | $l_{\text{mis}}$ (nm) | $d_{kT}$ (nm) | $\theta_{kT}$ (°) |
| Leu17 | K<sub>2</sub>L<sub>17</sub>K<sub>2</sub> | 2.55 | 3.3±0.5 | 23±10 | 2.3±0.4 | -1.0±0.9 | 3.2–3.3 | 4–45 |
| Leu19 | K<sub>2</sub>L<sub>19</sub>K<sub>2</sub> | 2.85 | 3.5±0.6 | 25±11 | 2.6±0.5 | -0.9±1.1 | 3.2–3.3 / 3.6–3.9 | 14–49 / 7–32 |
| Leu21 | K<sub>2</sub>L<sub>21</sub>K<sub>2</sub> | 3.15 | 3.8±0.6 | 26±12 | 2.8±0.6 | -1.0±1.2 | 3.7–4.0 / 4.3–4.4 | 8–39 / 11–25 |
| Leu23 | K<sub>2</sub>L<sub>23</sub>K<sub>2</sub> | 3.45 | 4.3±0.4 | 17±8 | 3.3±0.3 | -1.0±0.7 | 4.27–4.54 | 4–27 |
| Leu25 | K<sub>2</sub>L<sub>25</sub>K<sub>2</sub> | 3.75 | 4.5±0.4 | 23±9 | 3.4±0.5 | -1.1±0.9 | 4.32–4.56 | 10–36 |
| Leu27 | K<sub>2</sub>L<sub>27</sub>K<sub>2</sub> | 4.05 | 4.8±0.6 | 16±8 | 3.9±0.3 | -0.9±0.9 | 4.4–4.6 / 4.9–5.0 | 8–33 / 4–23 |
| Leu29 | K<sub>2</sub>L<sub>29</sub>K<sub>2</sub> | 4.35 | 5.1±0.2 | 23±8 | 4.0±0.5 | -1.1±0.7 | 4.32–4.55 | 27–48 |

We first studied whether the thickness gradient sorts the peptides, or whether alternative mechanisms—such as peptide tilt or membrane deformation—dominate. This is made possible by the thickness gradient, which—unlike biphasic setups ^20,21^—will induce a positiondependent lateral force on the peptides. The secondary structure of the peptides is fixed in the Martini 2.2 model,^18,19^ disallowing stretching, bending, or helix breaking, yet these are not expected to be important.^15,16^ We performed 3 sets of 100 μs-long unbiased simulations with varying initial positions of the peptides with a flat-bottom potential maintaining them within the linear regime of the thickness gradient (*x*=5–35 nm). As the peptides diffuse at rates of *D*≈0.015–0.027 nm^2^/ns, they cover 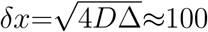 nm in the simulation time of Δ=100 μs, allowing them to spontaneously find their preferred membrane environment, while (large) MM likely renders this search faster. Rapid sorting indeed occurs as evidenced by the time traces of the peptide positions (Fig. S3 and movie at DOI: 10.6084/m9.figshare.24105606). We also repeated this calculation with 9 copies of Leu19, Leu23, or Leu27 present in the membrane and initially placed along the thickness gradient at constant intervals (Fig. S4 & DOI: 10.6084/m9.figshare.24105606). The TM peptide density profiles and tilt angle distributions are shown as colormaps in Fig. 2. Red markers show mean *±* standard deviation of the position of the peptide center of mass (and hence preferred thickness *d*_eq_, defined as inter-leaflet phosphate distance) or tilt angle (*θ*_eq_), which are also listed in Table 1. In one replica, Leu29 never left its initial position in the thin membrane (Fig. S3) and was hence omitted from the calculation of *d*_eq_ and *θ*_eq_.

**Figure 2:**
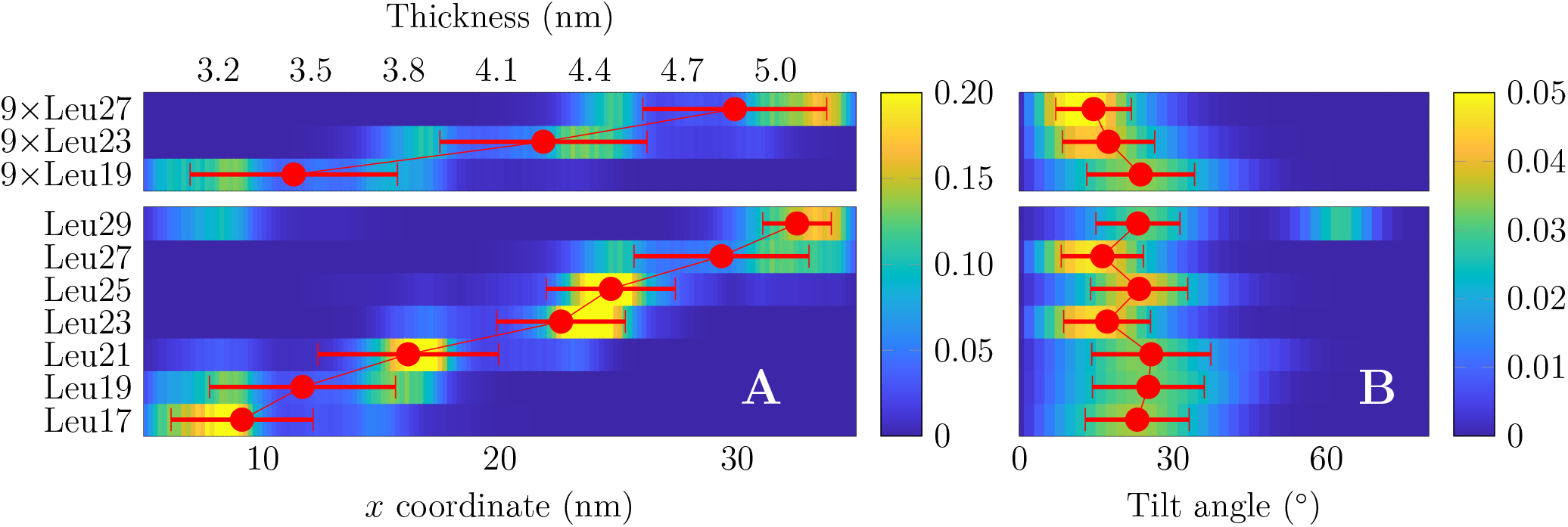
Spontaneous lateral sorting and tilting of the peptides. **A)** Density profiles of the peptides along the thickness gradient extracted from the last 50 μs of the unbiased simulations. Data shown for multi(top) and single-peptide (bottom) systems. For singlepeptide systems, data are gathered from 3 replicas with different initial peptide location. The mean value and standard deviation of the peptide center(s) of mass is shown in red markers. When initiated from a thin membrane, Leu29 did not find its equilibrium location and was omitted from the calculation of these values (Fig. S3). **B)** Histograms of the peptide tilt during the last 50 μs of the unbiased simulations. Red markers again show mean value and standard deviation with one replica for Leu29 omitted. All profiles integrate to 1, and the numeric values for red markers are listed in Table 1.

The *d*_eq_ values are 0.6–0.9 nm larger than the hydrophobic peptide lengths (*l*_TM_). Despite lateral sorting, the peptides still demonstrate a significant tilt *θ*_eq_ of 22*±*4^*°*^ on average without a systematic dependence on peptide length (Fig. 2B). Curiously, Leu23 and Leu27 tilt somewhat less (≈17^*°*^) than other peptides (≈24^*°*^), and this is reproduced in the multi-peptide system. Tilting leads to the projected peptide length *l*_proj_ along the *z* axis (normal to the membrane) being ≈0.3 nm shorter than *l*_TM_. Thus, the realized MM value (*l*_mis_=*d*_eq_–*l*_proj_) is consistently 1.0*±*0.1 nm for all studied peptides, whereas an extrapolation to zero lentgh provides a mismatch value of 0.8 nm (see TOC graphic). Concluding, spontaneous sorting takes place on the simulation time scale, while tilting also contributes to MM elimination.

To obtain physical insight into the sorting, we placed a single Leu17 or Leu29 at the center of the thickness gradient, and performed 100 independent unbiased simulations (Fig. 3A). We hypothesized that the velocity along the gradient (*x*) is linearly proportional to the distance of the peptide from its equilibrium location, 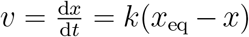, in other words proportional to MM. We solve for *x* = *x*_0_ − *A*(1 − exp(−*k/τ*)), which fits the data in Fig. 3A remarkably well (*R*^2^ *>* 0.99). This validates that *v*∼MM, which in viscous overdamped media where *F* ∼*v* also indicates that *F* ∼MM, *i*.*e*. a harmonic force. Curiously, both Leu17 (here MM*<*0) and Leu29 (MM*>*0) were found to relax towards *x*_eq_ with a time constant of 2.8 μs, suggesting that the force due to MM is symmetric around 0. Our peptides mainly respond to MM by tilting, yet membrane thickness deformation due to larger proteins was found to have a similar dependence on mismatch. ^25^

**Figure 3:**
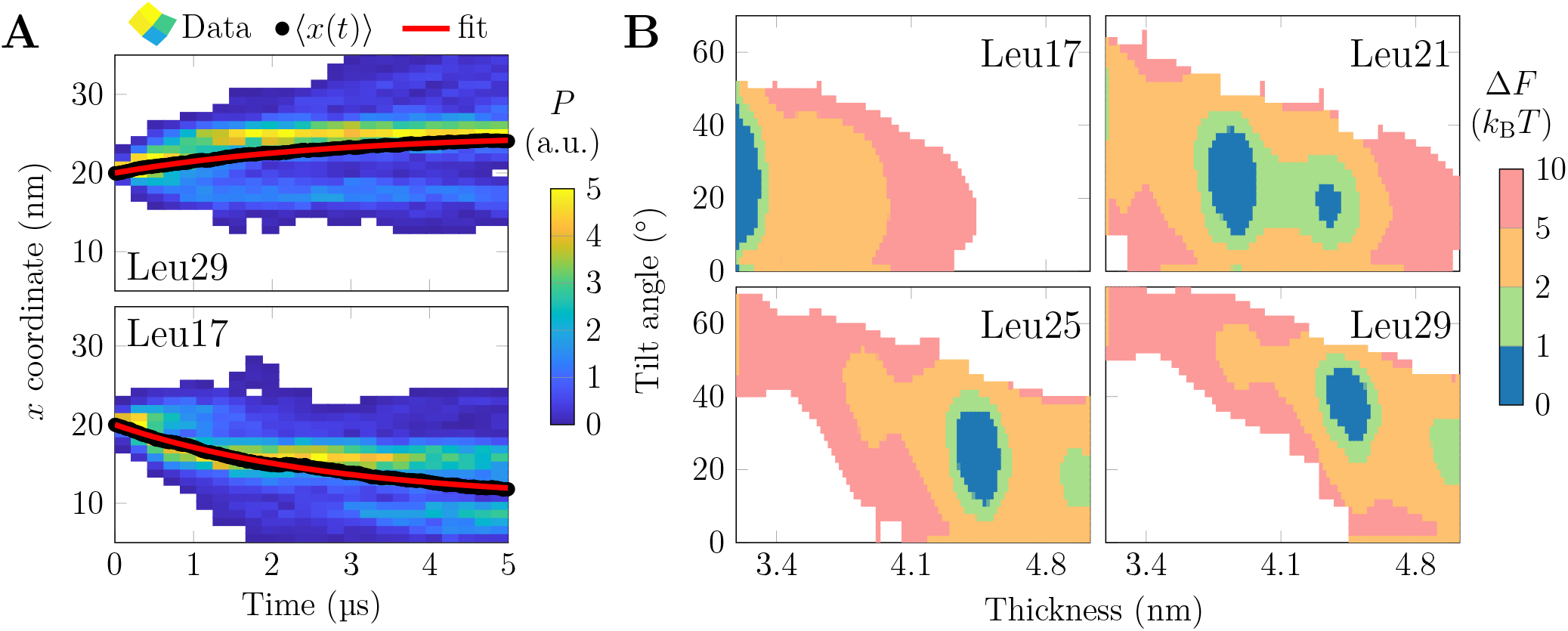
Energetics of lateral peptide sorting. **A:** Density of TM peptide center of mass as a function of time and *x* coordinate for the 100 replicas. Mean *x* shown in black and a fit of *x* = *x*_0_ − *A* (1− exp (−*t/τ*)) in red. **B:** 2D free energy surfaces as a function of membrane thickness and peptide tilt for selected systems. The rest are shown in Fig S5. Regions accessible from the global minimum within 1*×*, 2*×*, 5*×*, or 10 *× k*_B_*T* are highlighted, and Δ*F >* 10 *× k*_B_*T* discarded.

The peptides occasionally sample different thicknesses and tilt angles during the unbiased simulations (Fig. S3), indicating that the underlying free energy minima are shallow. To verify this, we performed umbrella sampling (US) simulations to extract potentials of mean force (PMFs) for the peptide position and hence as a function of local thickness (Δ*F* (*d*)). We combined this PMF with the free energy profile of peptide tilt angle obtained for each US window at as Δ*F* (*θ*)_tilt_ = −*k*_B_*T* ln (*P* (*θ*) */P* (*θ*_0_)) with *k*_B_*T* the thermal energy and *θ*_0_ the most likely tilt angle in that US window (Δ*F* (*θ*_0_) = 0). The sum Δ*F* (*d, θ*) ≈ Δ*F* (*d*) + Δ*F* (*θ*) was used to approximate the 2D free energy surface, and it provides both the minima as well as the thermally accessible ranges of thickness and tilt angle for each peptide (Selected ones in Fig. 3B, rest in Fig. S5). All profiles demonstrate minima which cover 0.1–0.3 nm in thickness (*d*_*kT*_) and 14–41^*°*^ in tilt angle (*θ*_*kT*_) within Δ*F* = *k*_B_*T* from the global minimum (ranges listed in Table 1; for peptides with two minima, values for both are reported). The profiles also reveal that doubling the threshold to 2 *× k*_B_*T* barely increases the accessible ranges, whereas within 5 *× k*_B_*T*, the peptides can already sample a thickness range *>* 1 nm. The accessible tilt angle range is less sensitive. Finally, within the 10 *× k*_B_*T* threshold, the peptides sample the entire available thickness range. The profiles demonstrate the expected trend; the thinner the membrane, the more tilted the peptides.

The entropic contribution leading to isotropic (polar) tilt angle *θ* dominates,^26^ yet the thickness gradient could also induce directional tilt. We applied the one-sample Kolmogorov– Smirnov test to estimate the *p* values for the hypothesis that the azimuth angle (*φ*) of the peptide differed from uniform distribution. The probability distribution of the *p* values in Fig. S6 demonstrates that in some cases the distribution deviates from the uniform one. However, these small *p* values seem to be distributed randomly among the peptides and among membrane thicknesses, indicating that there is no systematic tilt due to the gradient or the overlap regions inherent in our setup.

While the energetics of sorting by MM has not been previously studied, Kim and Im extracted the thermally accessible tilt angle ranges for WALP23/WALP27 peptides in POPC and DMPC membranes.^12^ WALP23/WALP27 have *l*_TM_ values similar to our Leu17/Leu21 peptides. We performed simulations of single-component POPC and DMPC membranes and identified the US windows with similar thicknesses. In the window corresponding to POPC, Leu17/Leu21 sample tilt angles of 5–29^*°*^/10–36^*°*^, whereas the values for WALP23/WALP27 in POPC were similar at 7–26^*°*^/14–46^*°*^.^12^ For DMPC-like thickness, Leu17/Leu21 showed tilts of 6–34^*°*^/18–47^*°*^, in reasonable agreement with WALP23/WALP27 in DMPC with 14–39^*°*^/32–51^*°*^.^12^ The small differences likely arise from different peptide sequences, especially the residues anchoring them to the membrane–water interfaces.

The US windows provide a systematic set of different (fixed) MM conditions. We first studied how the host membrane responds to the presence of the peptides of various lengths. We extracted the 2-dimensional thickness maps around for each peptide and for each US window. The thickness perturbations as a function of US window (peptide location) and membrane location are shown in Fig. S7. These maps demonstrate that the perturbed region spans a few nanometers around the peptide. We also extracted the values on the diagonal of these maps, *i*.*e*. the thickness perturbation values at the peptide position, and they are shown in Fig. 4A. These values clearly indicate that even the longest peptides have a meager (≈0.1 nm) thickening effect, whereas the shortest peptides render the thick membrane regions thinner by ≈0.5 nm). This asymmetry is perhaps not surprising, as the thickening is limited by the length of the acyl chains in their extended conformation, whereas thinning can occur to a larger extent *via* acyl chain tilt, disordering or even their interdigitation. This thinning contributes to MM estimates, but due to its complexity, we have omitted it from our other analyses. We next analyzed how the peptides respond to MM *via* tilting. Fig. 4A demonstrates tilt angles larger than 60^*°*^ in the case of significant positive MM. With thicker membranes, the tilt angles decrease monotonously until saturation at ≈10^*°*^. Our membrane does not contain a thick enough region to observe this saturation for the three longest peptides. We used these tilt values to calculate the projected peptide lengths (*l*_proj_) along the *z* axis (membrane normal), as for the unbiased simulations (Table 1). We then estimated the real MM defined as *l*_mis_=*d*_eq_–*l*_proj_ (Fig. 4B). Ideal MM (assuming peptide orientation along membrane normal) is also shown. In the case of positive ideal MM, peptide tilting leads to a fairly constant projected MM of –1 nm, in line with our unbiased simulations (Table 1). There is no obvious peptide length dependence. In the thicker membrane with ideal MM*<*–1 nm, the peptides can no longer tilt but rather maintain a constant tilt (Fig. 4A), and the projected MM follows the ideal behavior. Curiously, the magnitude of *l*_proj_ for each peptide is smallest right before it starts to follow the ideal MM scenario, *i*.*e*. at the least tilted conformation. This suggests that in too thin membranes, the peptides overcompensate for the MM by tilting.

**Figure 4:**
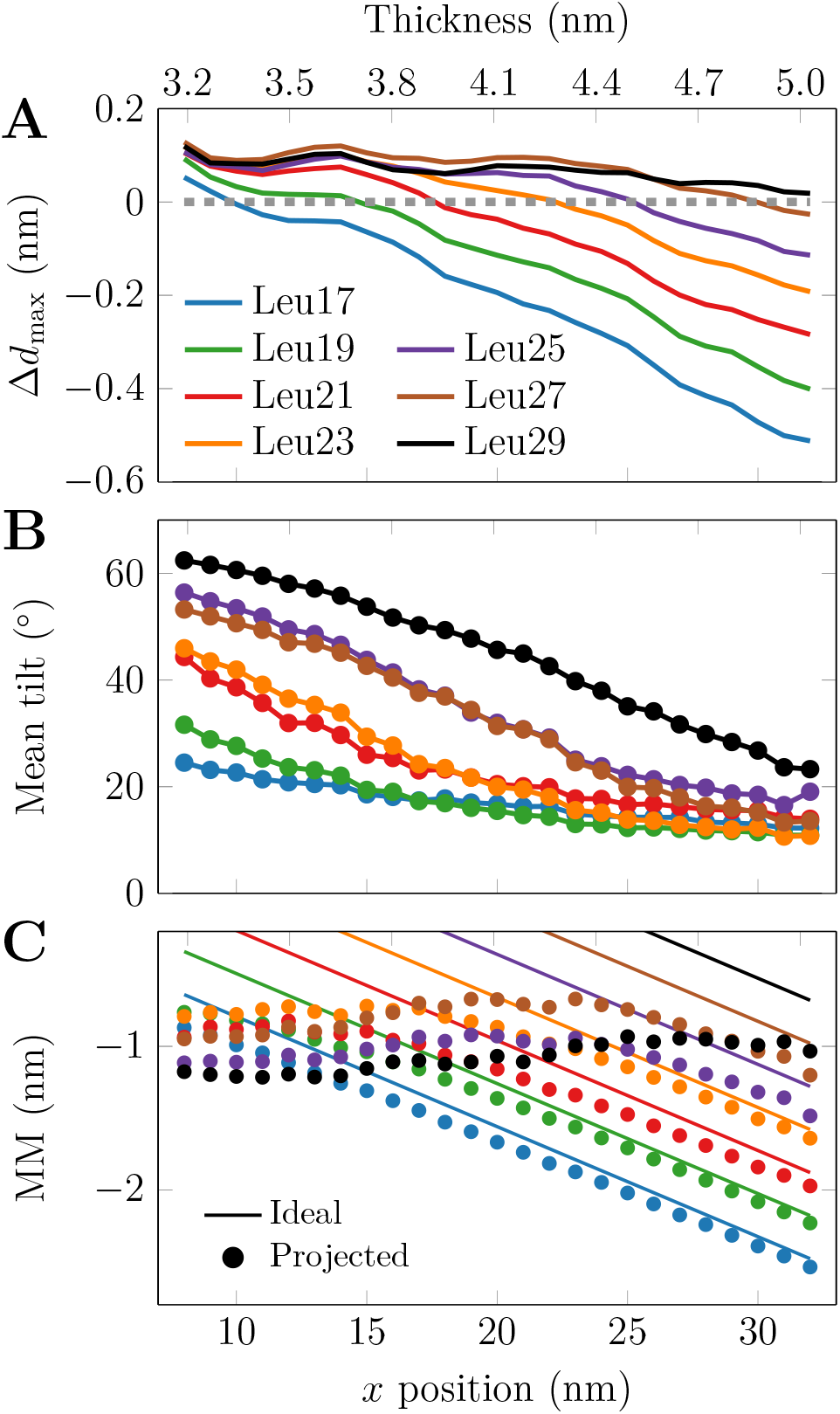
Peptide and membrane response to MM in US windows. **A)** Maximum perturbation of membrane thickness in the vicinity of the peptide. All peptides render the thin membrane regions slightly thicker by up to ≈ 0.1 nm, whereas the thinning effects are much more significant and up to ≈ 0.5 nm for the shortest peptide in the thin membrane regions. Data for the thickness perturbation in the entire membrane are available in Fig. S7. Peptide tilt as a function of thickness. Even at significant MM, the peptides maintain a tilt angle of 10–20^*°*^ due to entropy.^26^ **C)** Ideal and projected MMs. The peptides tilt to maintain a MM of –1 nm, yet at larger thicknesses the peptides adopt a small tilt angle and the projected MM follows the ideal MM.

Concluding, we have presented a novel simulation approach to study phenomena affected by membrane thickness. Using a combination of different lipid species and flat-bottom restraints, a thickness gradient is maintained along one axis of the simulation box. As the first example, we have focused on single-span peptides, which serve as model systems for the TMDs of physiologically important receptor tyrosine kinases. Our results demonstrate that peptides of different lengths are spontaneously sorted over distances of dozens of nanometers on the microsecond time scale. This indicates that our setup can be efficiently used to study the sorting of lipids, peptides, proteins, and other membrane-embedded objects. It can also be easily adapted to the study of larger membrane-spanning objects. However, for major protein complexes, the membrane dimensions might have to be extended, requiring the calibration of the overlap of the neighbouring phases, *i*.*e*. the adjustment of the widths of the flat-bottom potentials.

Moreover, with the *x* coordinate one-to-one mapped to thickness, conformational changes of proteins induced by the latter are readily studied. Free energy profiles of thickness-dependent properties—such as sorting or conformation—can be extracted using a simple reaction coordinate. We facilitate these and other yet unconsidered applications by providing all the simulation inputs and outputs in the Zenodo repository at DOIs: 10.5281/zenodo.10887673 & 10.5281/zenodo.10840054. Here, we used the CG Martini^18,19^ model, yet the future extension to atomistic resolution is straightforward. Morever, the CG approach has limited resolution, and although the smoothness of our thickness gradient is optimized (Fig. S2), there is still a small yet detectable bias of both the lipids (Fig. S1F) and the peptides (Fig. 3) towards the single-component membrane regions, rendering our approach only semi-quantitative. Other proposed approaches for the study of lipid sorting by mismatch seem to also suffer from boundary effects.^20–22^ In our setup, this is potentially an entropic effect, as a peptide in the overlap region decreases the total amount of lipid mixing permitted within the flat-bottom restraints. This is likely overcome with the additional resolution and hence smoother profiles of atomistic models.

## Acknowledgement

MJ thanks the Research Council of Finland (grant no. 338160) for funding, CSC–IT Center for Science (Espoo, Finland) for computational resources, and Dr. Tommi Kajander for fruitful discussions.

## Supporting Information Available

Details on simulation setup and analyses. Area per lipid of the membrane. Optimization of the overlap of flat-bottom potentials. Sorting of free lipids in the membrane. Sorting of peptides in unbiased simulations. All free energy profiles. Statistical tests on the randomness of the direction of peptide tilt. Membrane perturbation by the peptides.

