## Supplementary text and figures for "Energetics of the Transmembrane Peptide Sorting by Hydrophobic Mismatch"

### Supporting Information: Energetics of the Transmembrane Peptide Sorting by Hydrophobic Mismatch

#### System Setup

##### Lipid Membrane

The membrane with a thickness gradient was generated using four different lipid species with different acyl chains. In the Martini nomenclature, these are called DYPC (di-C12:1-C14:1 PC, 3 beads per acyl chain), DOPC (di-C16:1-C18:1 PC, 4 beads), DGPC (di-C20:1-C22:1 PC, 5 beads), and DNPC (di-C24:1-C26:1 PC, 6 beads). In each acyl chain, one bead corresponded to a hydrocarbon segment containing a double bond.

We generated patches for these lipid types separately using the `insane.py` tool.[S1] The DOPC and DGPC patches were 8×8 nm$^2$ in area, whereas the DYPC and DNPC ones were extended to 12×8 nm$^2$ as later justified. All boxes were 15 nm tall, fitting in enough water beads to ensure full hydration.

The patches were then catenated into a single elongated bilayer with an area of 40×8 nm$^2$. As DYPC and DNPC are located next to each other *via* the periodic boundary conditions, their mutual thickness mismatch leads to a perturbation at their interface. Thus, these patches were broader than the other two to ensure that there was also enough unperturbed DYPC and DNPC membrane present in the system.

Flat-bottom restraints acting along the $x$ axis, that is, direction of the thickness gradient were applied to the phosphate ("PO4") beads of each lipid. The center of the potential was always at the middle of the original patch, and its width was 2 nm broader than the original patch, thus leading to the mixing of lipids from neighbouring patches and a smoother thickness gradient. We also performed simulations with 0.5, 1.0, 2.0, 3.0, and 4.0 nm overlap.

The lipids-only systems were simulated for 5 µs, and they were primarily used to ensure that the setup provided a thickness gradient as smooth as possible considering the limited resolution of the coarse-grained approach.

#### Simulation With Free Lipids

To study the sorting of the different lipid types in the thickness gradient, we removed the flat-bottom restraints from 5 lipids of each lipid type in both leaflets. This system was simulated for 50 µs.

#### Transmembrane Peptides

We created polyleucine peptides ranging from 17 to 29 hydrophobic residues, in steps of 2 residues, using Avogadro.[S2] The peptides were capped at both ends by 2 lysine residues to anchor their termini to the membrane–water interfaces and thus ensure transmembrane (TM) orientation. All peptide sequences are given in Table 1 in the main text. The peptide structures were coarse-grained using the `martinize` tool. Next, a single copy of each peptide was inserted into the lipid membrane with the thickness gradient either to the center with average thickness ($x$=20 nm), or the region of thin ($x$=5 nm) or thick ($x$=35 nm) membrane.

To avoid the removal of any lipids during insertion, we simply merged the structure files of the peptide and the solvated lipid membrane, and energy-minimized the system to eliminate any steric contacts.

In addition, we embedded 9 copies of Leu19, Leu23, or Leu27 into the membrane with a thickness gradient at $x$ positions ranging form 7 nm to 31 nm with 3 nm intervals.

### Simulation Parameters

The simulation parameters are essentially identical for the lipids-only and the TM peptide-containing membranes. As the only difference, the protein was included in the same temperature coupling group as the lipids in the latter.

The flat-bottom potential that maintained the lipids at a certain range of the $x$ coordinate had a force constant of 100 kJ/mol/nm, and the width of the flat part was either 16 nm (DYPC and DNPC at the edges) or 12 nm (DOPC and DGPC in the middle) centered at the middle of the respective lipid patch. These values correspond to the originally generated single-component lipid patches, but extended to both directions along the $x$ axis by 2 nm for a smoother thickness gradient.

For all production runs, we used a time step of 25 fs. The lipid and peptide coordinates were written to the trajectories every 1 ns. Buffered Verlet lists[S3] were used to keep track of neighbor atoms. The reaction field electrostatics with a cut-off of 1.1 nm and $\epsilon_{\mathrm{rf}}$ of $\infty$ was used for efficiency.[S4] The charges were scaled by $\epsilon_r$ of 15. The van der Waals forces were cut off at 1.1 nm, and the potential was shifted to zero at this cutoff value. The stochastic velocity rescaling thermostat was used,[S5] and the temperatures of the membrane and the solvent were separately coupled to it. The target temperatures were set to 300 K and the time constants to 1 ps. The simulations were performed in the canonical ensemble, that is, under NVT conditions. Explicit constraints were handled by P-LINCS.[S6,S7]

For the systems containing TM peptides, we performed 100 µs-long unbiased simulations with three starting positions for each peptide. To avoid the peptides from partitioning outside of the desired [5 nm,35 nm] region containing the optimized thickness gradient (see Fig. 1B in the main text), we applied flat-bottom potential to all backbone beads of the peptides with a reference coordinate $x = 20$ nm, a flat regime with $r = 15$ nm and a force constant of 100 kJ/mol/nm$^2$ to keep them in the range [5 nm,35 nm].

In addition, we performed 100 independent runs for Leu17 and Leu29 in which the peptide was initially placed at intermediate thickness ($x$=20 nm). The peptide was first restrained during a 1 µs-long run, from which 100 independent configurations were extraxted (10 ns intervals). Simulations initiated from these 100 configurations were 5 µs-long each.

Additionally, we performed umbrella sampling (US) simulations, where the reaction coordinate was defined by the position of the peptide center of mass along the $x$ axis (*i.e.* along the thickness gradient). An absolute reference point at $x = 20$ nm was used. The umbrella windows covered the $x$ coordinate range from 8 nm to 32 nm with 49 windows separated by 0.5 nm. Every other window (at 8 nm, 9 nm, ...) was simulated for 50 µs, whereas the rest (at 8.5 nm, 9.5 nm, ...) were simulated for 10 µs each.

In terms of implementation, we set the reaction coordinate geometry into "direction", the respective vector into [1 0 0] (pulling only along the $x$ axis, and sampled the reaction coordinates in the range of [-12,12]. The force constant was set to 100 kJ/mol/nm$^2$, which resulted in adequate overlap of the position distributions for neighbouring windows.

The total simulation time was 13.5 ms, consisting of 3.1 ms of unbiased simulations and 10.4 ms of US simulations. The simulation input parameter files, as well as all other input and output files for the spontaneous sorting simulations as well as the umbrella sampling windows, are available in the Zenodo repository at DOIs: 10.5281/zenodo.10887673 & 10.5281/zenodo.10840054.

### Analysis Methods

From the simulation of the lipid membrane alone, we used `g_lomepro`[S8] to extract the thickness profile based on the phosphate ("PO4") bead locations. The `gmx density` tool bundled with GROMACS was used to extract lipid density, which was then converted to area per lipid.

The simulations with free lipids was analyzed using the `gmx density` tool. The density along the thickness gradient ($x$) was converted to a free energy profile by $\Delta G = -RT \ln(\rho/\rho_0)$, where $\rho_0$ is the density at the most likely location along the gradient.

From the unbiased simulations with the peptides, their center of mass $x$ coordinates and peptide densities were extracted using `gmx traj` and `gmx density` of GROMACS, respectively. The peptide tilt was calculated using `gmx gangle`. As the peptide structure is rigid in Martini, the backbone beads ("BB") of the first and last non-lysine residue were used to define the helix orientation. The last 50 µs of the trajectories were used for the densities and the position and tilt distributions.

From the 100 unbiased short simulations with Leu17 and Leu29, we extracted the $x$ coordinate of the peptide center of mass using `gmx traj`. These coordinates were used to demonstrate that the peptides obtain a velocity directly proportional to the hydrophobic mismatch.

From the US simulations with the peptides, the potential of mean force (PMF) along the $x$ coodinate, *i.e.* along the thickness gradient was extracted using the `gmx wham` tool of GROMACS. The first 10 ns of the simulation output were discarded from the analysis.

The membrane perturbation was analyzed from the US windows by first extracting the 2-dimensional thickness maps with respect to the peptide position and with a grid spacing of 0.25 nm. A slab with a width of 2 nm along the $y$ axis and with the protein at its center was analyzed using `g_lomepro`.[S8] The data were averaged over this range of $y$ coordinates. The reference thickness profile (in the absence of peptides) along the $x$ coordinate was subtracted from these data, and the result plotted for each peptide and each US window in a

2-dimensional map. In addition, the values from the diagonal of these maps, corresponding to the perturbation at the immediate vicinity of the peptide were also extracted.

To investigate whether the axis of rotation of the peptides exhibits a a systematic tilt, we performed one-sample Kolmogorov–Smirnov tests (as implemented in SciPy[S9]) of the azimuthal angle, $\phi$, of the vector connecting the N-terminal and C-terminal backbone beads against a uniform distribution. We sub-sampled the trajectory and used every 40$^{\text{th}}$ frame to decrease correlations in the $\phi$ angles while still retaining enough data for statistical averaging.

### Additional Results

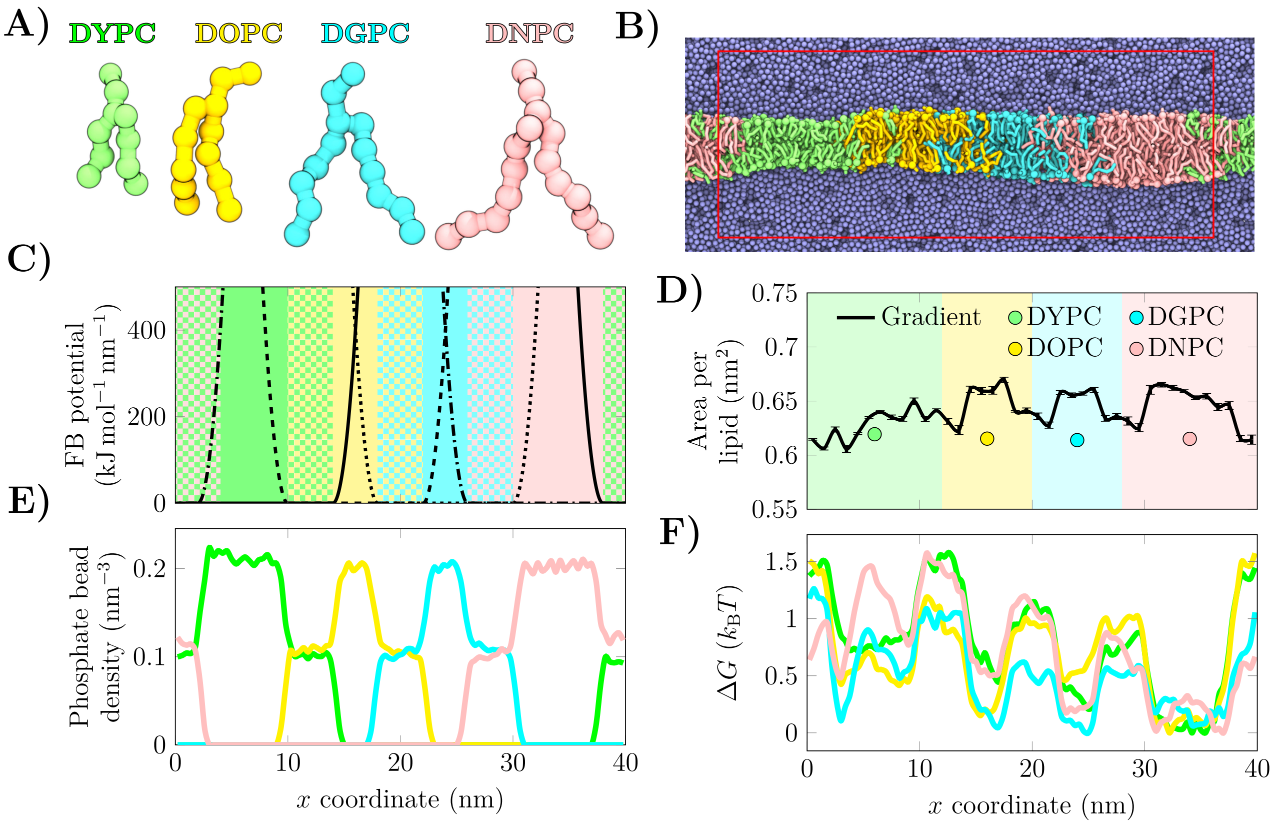


Figure S1: Properties of the membrane in the absence of peptides. **A)** Structures of the used lipids. **B)** Snapshot of the simulation setup. The red rectangle denotes the simulation cell. **C)** The shapes of the flat-bottom restraints ("FB potential"). The FB potentials for the different lipid types are shown in solid (DYPC), dashed (DOPC), dotted (DGPC) or dot-dashed (DNPC) lines. The single-lipid patches are colored with a solid color, whereas those with overlap are shown with the corresponding checkered patterns. **D)** Area per lipid as a function of $x$ coordinate in the gradient membrane. Values extracted from both peptide-free simulations (solid line) and from single-component membranes (coloured markers) are shown for comparison. **E)** Densities of the phosphate beads of different lipid types along the thickness gradient. **F)** Free energy profile for the free lipids. The sorting tendency for lipids is minor at $< 1\ k_{\mathrm{B}}T$. Despite the optimal overlap (see Fig. S2 and Fig. 1 in the main text), the mixed regions induce a small repulsive bias of $\approx 1\ k_{\mathrm{B}}T$.

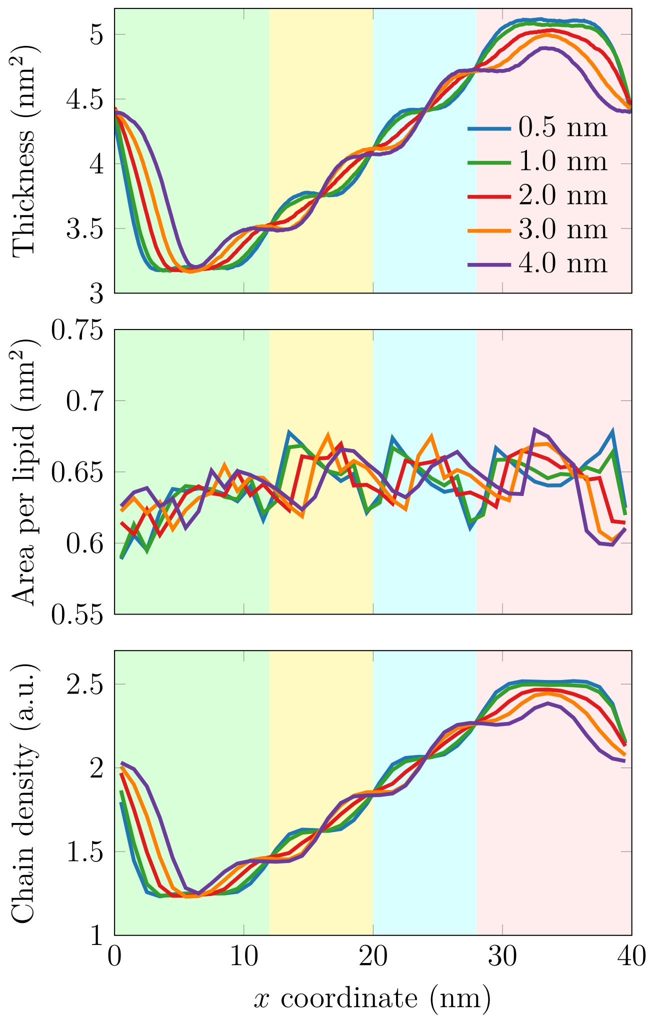


Figure S2: Effect of overlap of the flat-bottom restraints maintaining the thickness gradient. For different amounts of overlap, thickness, area per lipid, and chain density profiles across $x$ axis, *i.e.* along the thickness gradient are shown. For overlaps other than 2 nm, a step-wise pattern emerges in the thickness and chain density profiles. In area per lipid, the overlap has little effect on the profile. This suggests that an overlap of 2 nm, used in the rest of the simulations in this work, leads to the smallest deviation from a linear thickness gradient without inducing artefacts in area per lipid nor chain density. Shading is as in Fig. S1.

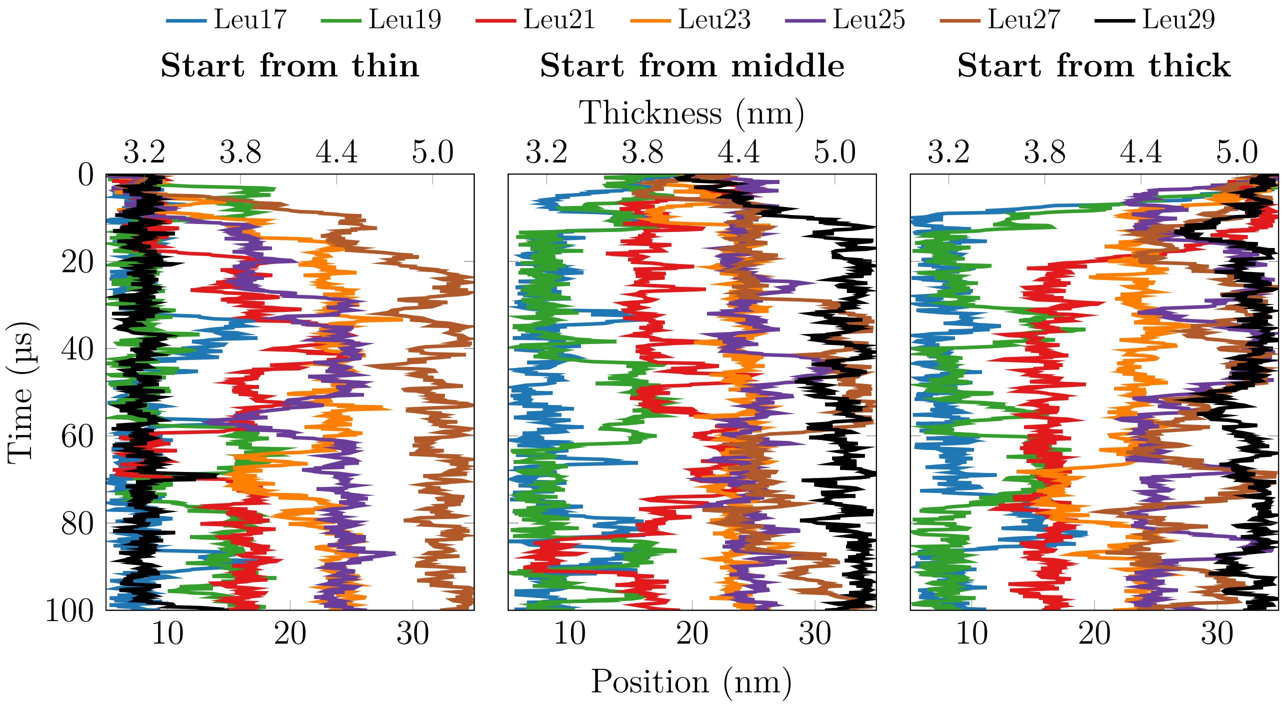


Figure S3: Sorting of the peptides in the unbiased single-peptide simulations with three initial conditions.

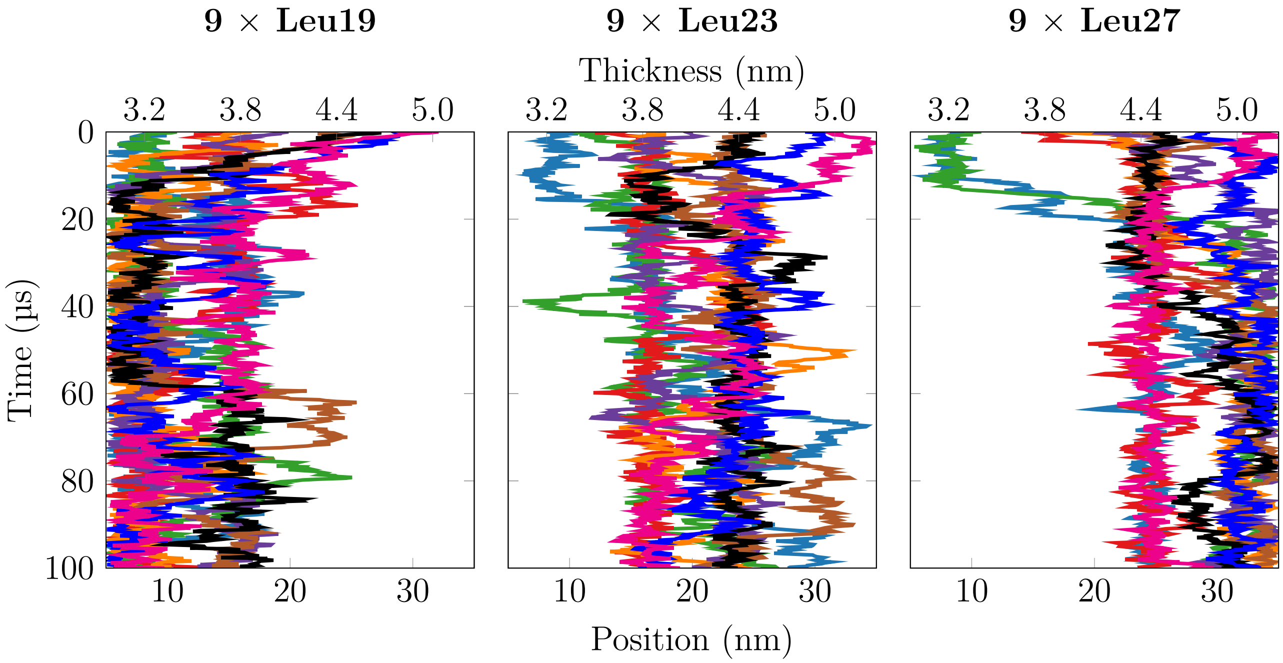


Figure S4: Sorting of the peptides in unbiased simulations with multiple peptides present in the membrane. Each peptide is shown with a different color.

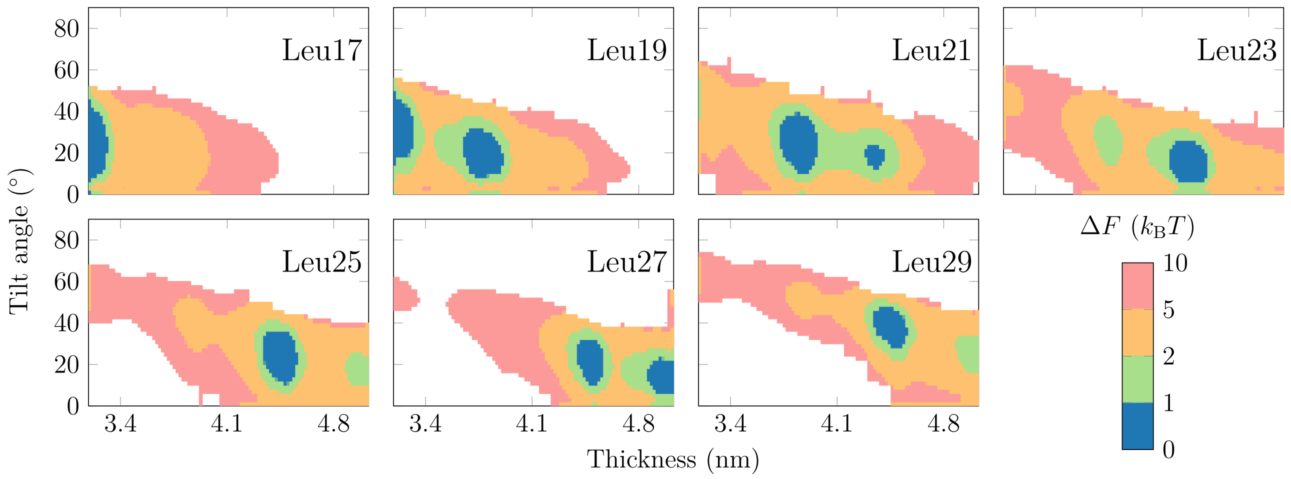


Figure S5: 2D free energy surfaces as a function of membrane thickness and peptide tilt for all systems. Selected ones are shown in Fig. 3 in the main text.

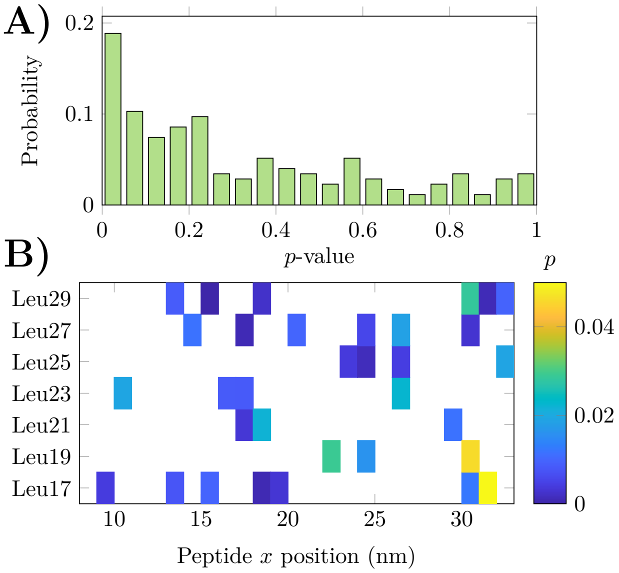


Figure S6: Results from the Kolmogorov–Smirnov test. **A)** Total distribution of $p$ values extracted for all peptides and all US windows. **B)** The distribution of US windows for which $p < 0.05$.

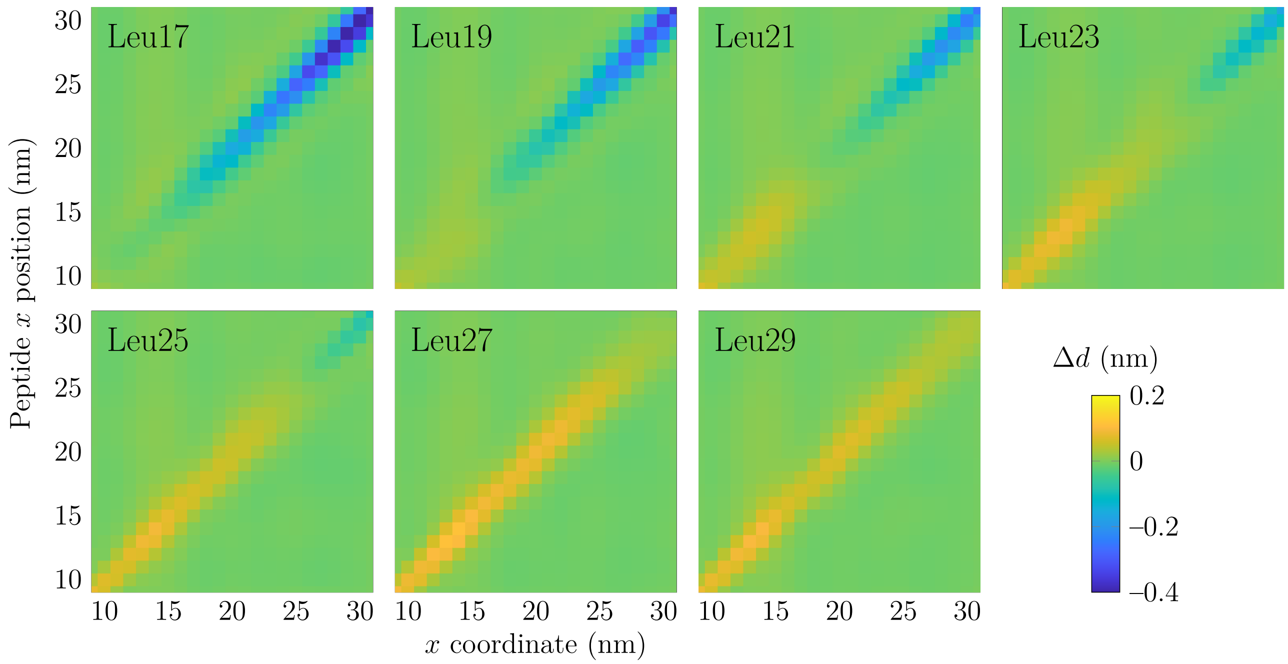


Figure S7: Membrane thickness perturbation by the peptides. The rows ("Peptide $x$ position") show data for the different US windows (*i.e.* the restrained position of the peptide), whereas the columns ("$x$ coordinate") show the different membrane locations. Unsurprisingly, the perturbation is localized at near the peptide, and extends up to a few nanometers along $x$. The values on the diagonal (*i.e.* the perturbation at the peptide location) are shown in Fig. 4A in the main text.
